# Organizing principles of whole-brain functional connectivity in zebrafish larvae

**DOI:** 10.1101/496414

**Authors:** Richard F. Betzel

## Abstract

Network science has begun to reveal the fundamental principles by which large-scale brain networks are organized, including geometric constraints, a balance between segregative and integrative features, and flexible brain areas. However, it remains unknown whether whole-brain networks imaged at the cellular level are organized according to similar principles. Here, we analyze whole-brain functional networks reconstructed from calcium imaging data recorded in larval zebrafish. Our analyses reveal that functional connections are distance-dependent and that networks exhibit hierarchical modular structure and hubs that span module boundaries. We go on to show that spontaneous network structure places constraints on stimulus-evoked reconfigurations of connections and that networks are highly consistent across individuals. Our analyses reveal, for the first time, basic organizing principles of whole-brain functional brain networks at the microscale. Our overarching methodological framework provides a blueprint for studying the correlated activity at the cellular level using a low-dimensional network representation. Our work forms a conceptual bridge between macro- and microscale network neuroscience and opens myriad paths for future studies to investigate network structure of nervous systems at the cellular level.

## INTRODUCTION

Nervous systems are collections of functionally and structurally connected neurons, neuronal populations, and brain areas [1]. Coordination of and within these networks underpins an organism’s ability to process sensory stimuli [2, 3], successfully navigate its environment [4, 5], and to perform goal-directed action [6]. Network science provides a quantitative framework for representing and analyzing the organization of biological neural networks [7]. Within this framework, neural elements and their pairwise interactions are modeled as the nodes and edges of a complex network [8], to which one can apply a growing suite of powerful graph-theoretic tools to assay the network’s structural [9], functional [10], and dynamical [11] properties.

Though the network model can be applied to nervous systems imaged at virtually any spatial scale [12–14], the majority of applications thus far have focused on the macroscale, where nodes represent brain regions and edges represent pairwise statistical associations of recorded activity (functional connectivity; FC). Though macroscale network analyses have linked variation in network features to cognition [10], disease [15], and development [16], they have also begun to elucidate the general principles by which biological neural networks are organized [17, 18].

These principles include a balance between network structures that support segregative (local and specialized) and integrative (global and generalized) information processing [19, 20], e.g. modules *versus* hubs [21–24], a strong brain-wide drive to reduce the material and metabolic cost of wiring [25–27], and an intrinsic functional architecture that reconfigures subtly and efficiently in response to tasks or stimulation [28, 29].

While these organizing principles have been observed at the macroscale across individuals and phylogeny [30, 31], it remains unclear whether analogous principles shape the architecture of nervous systems at the microscale, where networks represent interactions among cells and molecules [13, 32]. Though the number of studies investigating microscale network structure continues to grow [33–38], an emphasis on neuronal populations as the unit of interest (rather than the brain as a whole) has prevented the uncovering of the general principles by which microscale networks are organized.

Currently, the organization principles underpinning whole-brain microscale networks remains largely unexplored. Here, we take advantage of recent technological advances to investigate spontaneous and stimulus-evoked FC in zebrafish larvae [39–42]. Our analyses reveal several putative organizing principles. These include strong geometric constraints on the magnitude and valence of connection weights, evidence of hierarchical and multi-scale modular structure, balanced by the presence of polyfunctional hubs. We show that spontaneous and stimulus-evoked networks are highly similar. Nonetheless, we also find evidence of stimulus-driven module re-configuration. Interestingly, the nodes with the greatest propensity for reconfiguration overlap with the same polyfunctional hubs uncovered under spontaneous conditions, linking intrinsic network architecture to behavior. In summary, our findings link whole-brain macro- and micro-scale analyses and highlight network science as a framework for bridging neuroscientific inquiry across spatial scales and domains.

## RESULTS

Here, we aimed to uncover organizing principles of spontaneous and stimulus-evoked FC in zebrafish larvae. All details concerning data acquisition and network construction are included in the **Materials and Methods** section. The following subsections are organized as follows. First, we analyze group-representative spontaneous FC to identify signatures of geometric constraints, hierarchical modular structure, and polyfunctional hubs. Next, we extend these analyses to stimulus-evoked FC and show that stimulus-evoked FC is constrained by the brain’s intrinsic (spontaneous) functional organization. Finally, we present results of single-subject analyses showing a high-level of inter-subject consistency.

### Geometric organization of spontaneous FC

Previous analyses of whole-brain FC have revealed that connection weights and other network features are shaped by underlying geometric relationships, e.g. nodes’ locations in space [43–45]. These constraints result in networks that favor strong, short-range connections and are believed to reflect brain-wide drives to reduce the metabolic cost of coordinating activity between brain areas over long distances [46]. However, it remains largely unknown whether whole-brain microscale networks are subject to similar constraints. To address this question, we analyzed single-cell calcium fluorescence traces from zebrafish larvae recorded in stimulus-free (i.e. spontaneous) conditions. We aggregated cells into *N* = 256 hemispherically-symmetric, functionally-homogeneous, and spatially-localized parcels (nodes; Fig. 1a; see **Materials and Methods** for details), calculated the average fluorescence trace for each node, and computed *A* = {*A_ij_*}, the full matrix of Fisher-transformed Pearson correlations (Fig. 1b; see **Materials and Methods** for preprocessing details). Unless otherwise noted, all subsequent analyses were carried out on this matrix *A*, which we regarded as a fully-weighted and signed connectivity matrix.

**FIG. 1.**
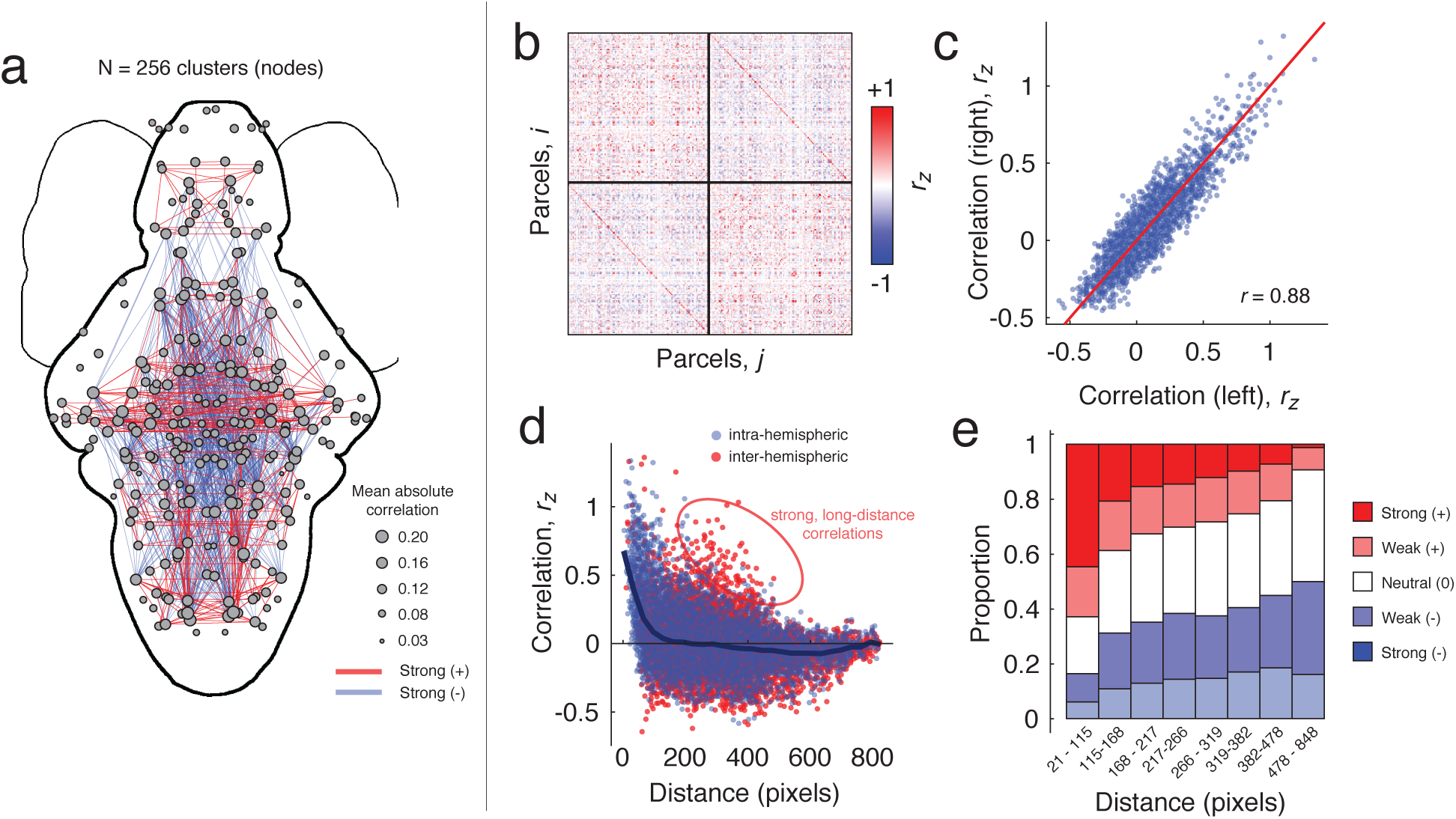
Spatial features of zebrafish whole-brain functional connectivity. (*a*) Thresholded network in anatomical space. Nodes represent parcels with size proportional to average absolute correlation. Red and blue lines represent top 400 positive and negative correlations/anti-correlations according to magnitude. (*b*) Correlation matrix, *A*, ordered by hemisphere. (*c*) Scatterplot of left *versus* right within-hemisphere connectivity (*d*) Scatterplot of straight-line distance between parcel centroids and their correlation magnitude. Within- and between-hemisphere connections are plotted separately. Note that in both cases, connection weight decays as a function of distance. However, there nonetheless exists a small fraction of long-distance inter-hemispheric connections. (*e*) Breakdown of connection types by distance bins. Note that in general, the proportion of positive (strong or weak) correlations decreases with distance, while the prevalence of anti-correlations increases with over longer distances.

First, to assess whether FC exhibited hemispheric symmetries, we calculated the similarity of all left and right within-hemisphere connections (Fig. 1c). We observed that within-hemisphere connectivity patterns were highly correlated (*r* = 0.88; *p* < 0.05). To further assess the relationship of network architecture with geometry, we then plotted connection weight as a function of the Euclidean distance between parcel centroids. We found that, on average, connection weight decayed monotonically as a function of distance. However, we also observed a small subset of inter-hemispheric connections that were unexpectedly strong given their length (Fig. 1d). A similar pattern was observed when we classified connections according to their valence and magnitude, and examined the proportion of each class within a fixed set of distance bins (Fig. 1e). In general, nodes separated by short distances tended to be linked by strong, positive correlations. At longer distances, however, the proportion of positive correlations decreased and was overtaken by an increase in neutral to strong anti-correlations.

These observations confirm that geometric relationships serve as powerful determinants of connections’ strengths and valences. Despite the fact that connection weight (on average) decreases monotonically with distance, the presence of strong, long-distance correlations suggests that geometry insufficiently explains brain-wide patterns of FC, and that coordination of activity over long distances may act as an additional functional constraint on network architecture. Collectively, these findings are analogous to those observed at the macroscale [43, 45] and draw a clear conceptual link between the organization of biological neural networks at the macro- and microscales.

### Modular organization of spontaneous FC

In the previous section, and in agreement with observations made at the macroscale, we suggested that a monotonically decaying relationship of connection weight with distance may serve as a key organizing principle responsible for shaping the architecture of biological neural networks. Here and in the next section, we explore another putative organizing principle. Namely, the requirement that biological neural networks balance features that support both segregated (localized) and integrated (global) brain function, i.e. network modules and hubs, respectively [19, 20].

Modules are groups of nodes that are densely connected to one another, but weakly connected between groups [21, 22]. Because of modules’ near-autonomy from one another, they are thought to represent groups of nodes that perform the same or similar brain function and are believed to engender specialized information processing. In general, modules are not restricted to a single topological scale and can be arranged hierarchically, with deep levels of the hierarchy reflecting increasing functional specialization [12, 47, 48].

While modular organization has been frequently observed in whole-brain macroscale networks [21, 49, 50], little is known about the modular structure of whole-brain networks at the microscale [37, 38, 51]. Here, we leverage recent advances in modularity maximization, a data-driven method for uncovering a network’s modules, to uncover the hierarchical modular structure of spontaneous FC [52] (see **Materials and Methods** for more details).

Modularity maximization returns hierarchically-related partitions of nodes into modules, where the hierarchical levels are determined using a statistical criterion [52]. We found evidence of a hierarchy comprising 24 distinct levels, with the number of modules at any level ranging from 2 to 27 (Fig. 2a). For the sake of brevity, we show partitions of nodes into *c* = 2 (Fig. 2a-c), 4 (Fig. 2d-f), and 9 (Fig. 2g-i) modules.

**FIG. 2.**
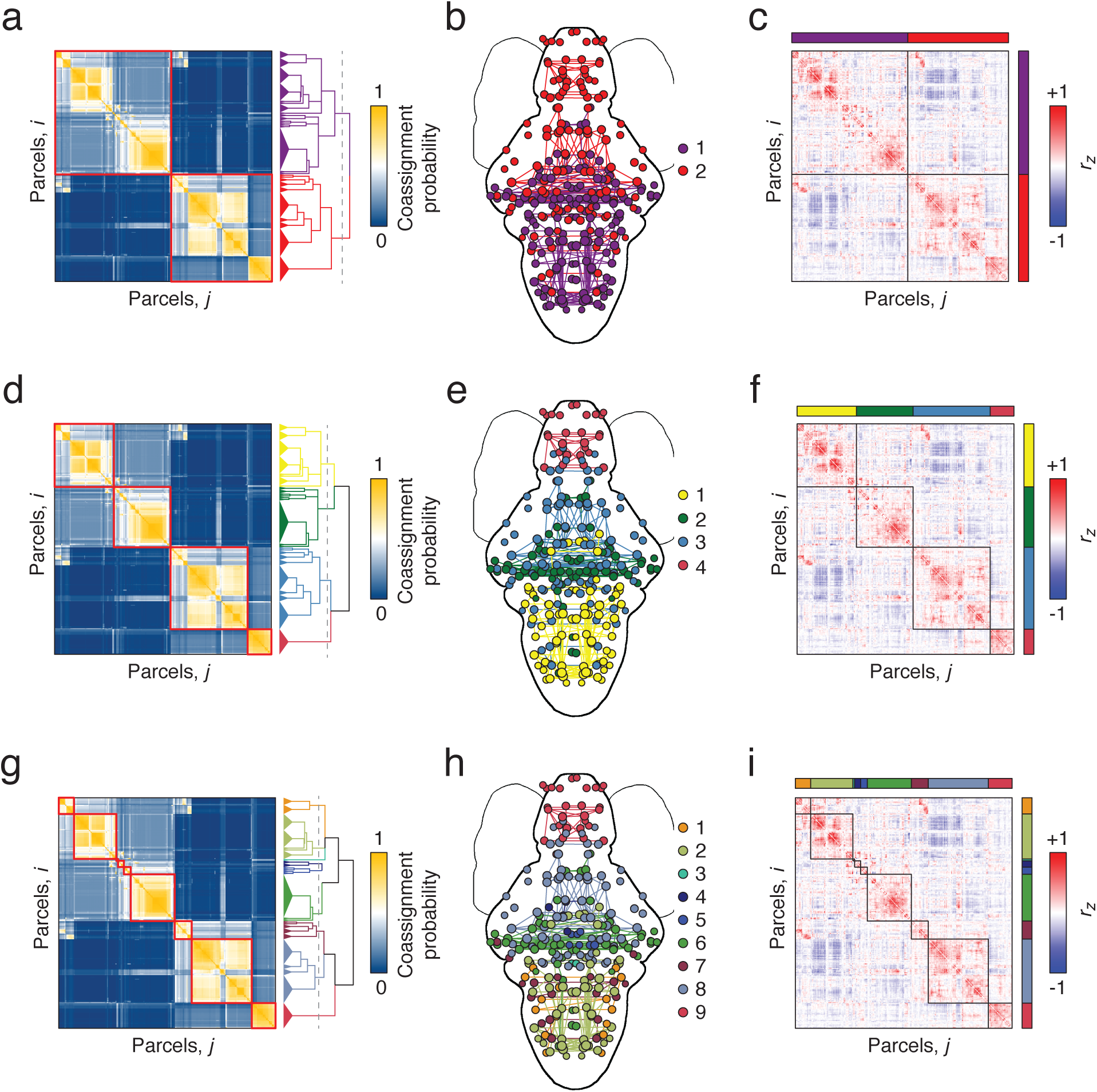
Hierarchical and multi-resolution modular structure. (*a*) Co-assignment matrix with with hierarchy cut to highlight a two-module division (red blocks). (*b*) Modules plotted in anatomical space. Nodes are colored according to their module assignment, and within-module connections are colored similarly. (*c*) Correlation matrix ordered and blocked to highlight modular structure. Panels *d*-*f* and *g*-*i* show similar figures but with the number of modules equal to four and nine, respectively.

Many of the modules recapitulate known functional and anatomical divisions of zebrafish. For instance, the red module (labeled 4 in Fig. 2e and 9 in Fig. 2h) circumscribes the telencephalnon. Interestingly, this module isolates itself early within the hierarchy, and exhibits no clear sub-divisions at deeper levels. Similarly, module 2 in Fig. 2e and modules 4, 5, 6 and 8 in Fig. 2h represent coarse and fine divisions of the mesencephalon. Unlike the telencephalonic cluster, we identify multiple mesencephalonic subdivisions, suggesting that these areas have the capacity to subtend varying levels of functional specialization.

Collectively, these findings suggest that the spontaneous FC is organized hierarchically into modules that exhibit clear mappings to known neuroanatomy. These findings are in close agreement with analogous investigations of whole-brain networks at the macroscale [22] and suggest that modular organization may be a unifying principle by which brain networks at all scales are structured [53].

### Hub organization of spontaneous FC

In the previous section we presented evidence that zebrafish spontaneous FC exhibits hierarchical modular structure. While segregated modules may be useful for the development of specialized brain function, complex behavior also requires network features that support the integration of information across different modules [20, 54, 55]. One class of network feature that supports precisely this type of processing is hubs, nodes whose connections straddle the boundaries of modules. Here, we identify hubs in spontaneous FC using the network measure *participation coefficient* [56].

Participation coefficient measures the uniformity with which a node’s connections are distributed across modules; values close to 1 or 0 indicate nodes that connect to many different modules or are concentrated within a small number of modules, respectively. We illustrate this concept schematically in Fig. 3a. where we show examples of two nodes – one with low (top) and another with high (bottom) participation. Given the modules detected in the previous section, we calculated each node’s participation coefficient and averaged participation scores over all partitions composed of the same number of modules. We focus, here, on partitions of the network into 2 - 25 modules (Fig. 3b). Averaging participation coefficient across this range, we find marked heterogeneity in the spatial locations of high-participation hubs, with the greatest concentration appearing in the hindbrain within the rhombencephalon (Fig. 3c). Though brain-wide patterns of participation coefficient are largely stable as we vary the number of modules, we nonetheless observe subtle scale dependencies (Fig. 3d), suggesting that small subsets of nodes may be uniquely configured to act as hubs at one particular scale, but less so at another.

**FIG. 3.**
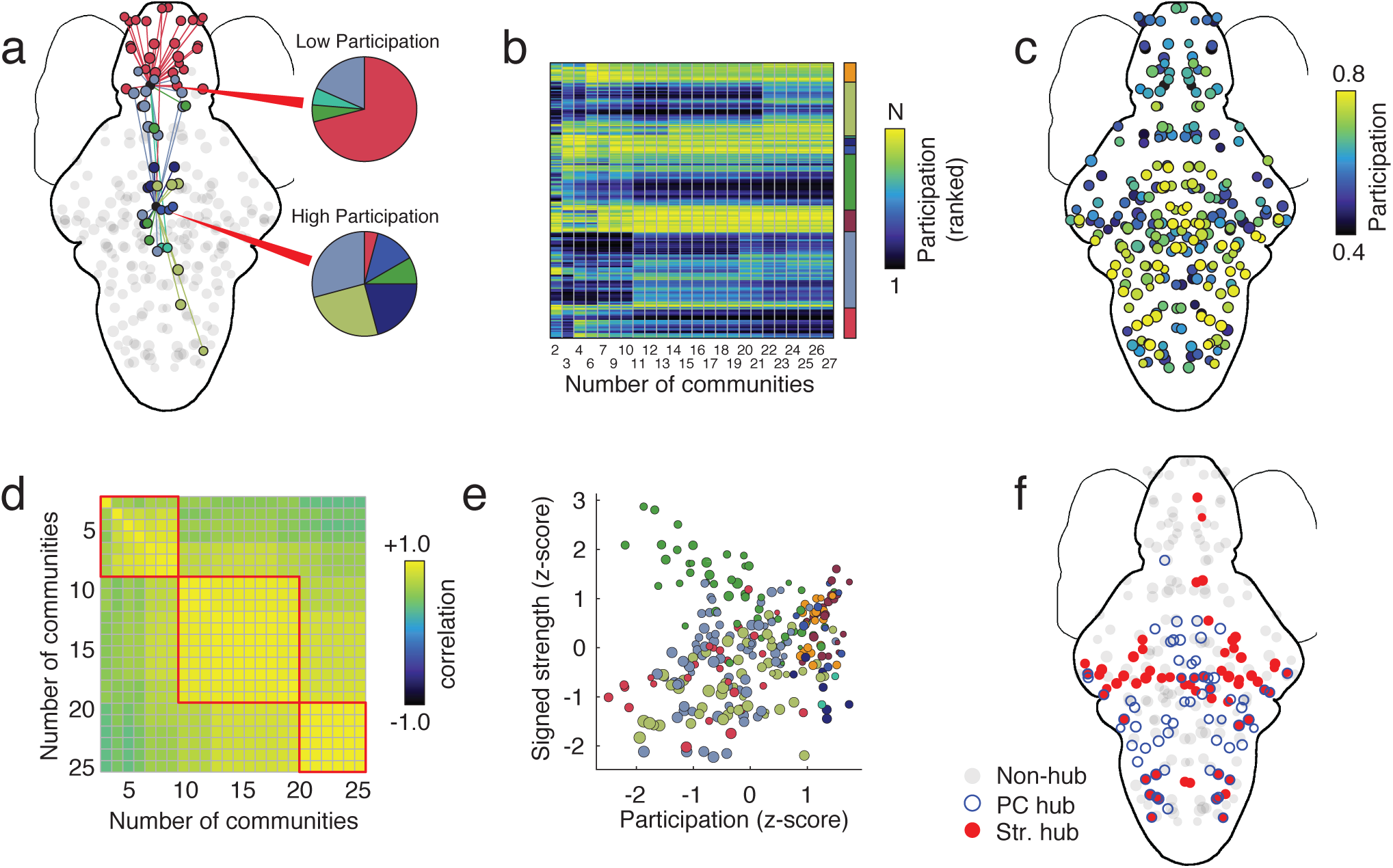
Participation coefficient and hub classification. (*a*) Schematic illustration of low- and high-participation nodes. Connections made by nodes with low participation (non-hubs) are concentrated within those nodes’ modules; connections made by nodes with high participation (hubs) are distributed across many modules. (*b*) Mean participation coefficient (ranked) as a function of the number of detected modules. (*c*) Average participation coefficient when the number of modules ranges from 2 to 25. (*d*) Correlation of brain-wide participation coefficient maps for different numbers of modules. While the changes in participation coefficient maps are subtle, there nonetheless appear to be three “regimes” corresponding to distinct patterns of participation. (*e*) Participation coefficient and signed strength, z-scored and plotted against one another. Both measures serve as indices of “hubness” but are, on average, uncorrelated. (*f*) Nodes in the top 25^th^ percentile by participation and signed strength. On average, there is little overlap.

To better contextualize these findings, we combined nodes’ participation coefficients with their signed strength (total weight of functional connections), another measure sometimes used for identifying hubs (Fig. 3e). In general, we find that these two metrics are uncorrelated, i.e. the most hub-like nodes according to participation coefficient and strength are not necessarily overlapping. Indeed, when we examine the top 25% of hubs according to each measure, we find little overlap between the two hub measures (Fig. 3f).

These findings, combined with those from the previous section, suggest that spontaneous FC at the microscale strikes a precarious balance between features that promote segregated (i.e. local) brain function and those that promote integrative (i.e. global) brain function. These findings mirror those reported in macroscale networks [20], where the expression of hub areas has been linked to both genetics and cognitive performance [57].

### Correspondence of spontaneous and stimulus-evoked FC

In the previous sections, we focused on organizing principles of spontaneous (i.e. stimulus-free) FC. Spontaneous FC represents an intrinsic, baseline state in which functional connections are shaped primarily by an underlying network of physical pathways, projections, and fiber tracts rather than the functional demands of a task or stimulus [10, 58, 59]. Nonetheless, nervous systems must be flexible and capable of reconfiguring in order meet those demands should they arise [28, 60]. What features characterize stimulus-induced reconfiguration? Is it a wholesale reorganization of the network? Is it restricted to a small subset of connections or nodes? In this section, we explore the effect of different stimuli on FC organization.

To study stimulus-evoked changes in functional network organization, we estimated FC during the presentation of different visual stimuli. In addition to spontaneous activity, we considered: Phototaxis (PT), Optomotor response (OMR), Looming response (Looming), and Dark-flash response (DF) (See **Materials and Methods** for more details).

First we estimated a FC matrix separately for each stimulus condition (Fig. 4a). To assess the similarity of these matrices, we extracted their upper triangle elements, computed pairwise inter-stimulus correlations (Fig. 4b), and also generated a scatterplot of these elements against one another (Fig. 4c). Overall, we found that the stimulus-evoked connectivity matrices were highly similar (mean correlation of *r* = 0.72; *p* < 0.05, Bonferroni corrected). These results support the hypothesis that the differences between spontaneous and stimulus-evoked connectivity is characterized by subtle shifts in connection weights and not by a wholesale reorganization of FC. In the next section, we explore these subtleties in greater detail.

**FIG. 4.**
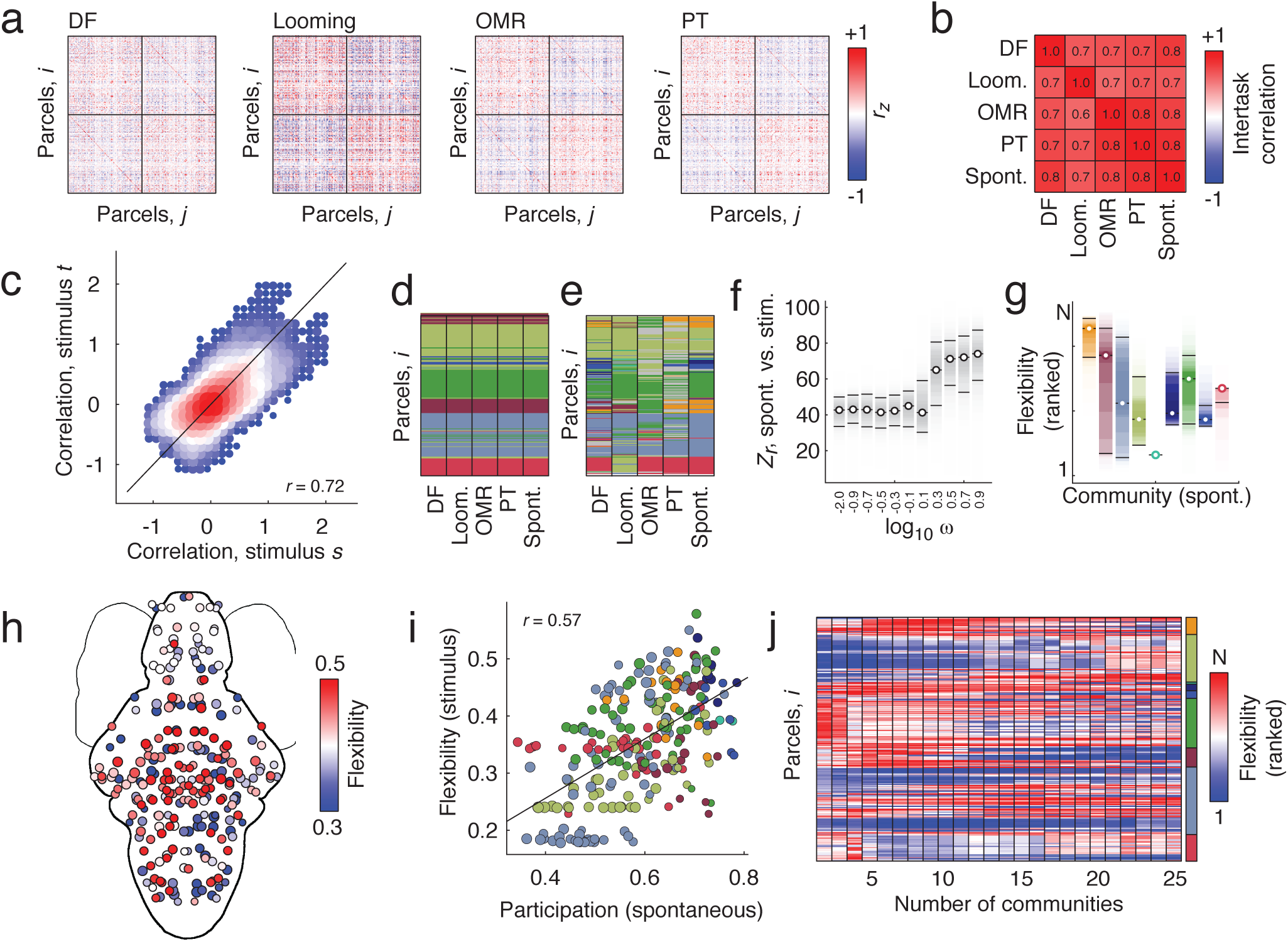
Comparing spontaneous and stimulus-evoked network architecture. (*a*) Functional connectivity matrices for four different stimulus conditions: Dark-flash (DF), Looming, optomotor response (OMR), and phototaxis (PT). (*b*) Upper triangle correlation for four stimulus conditions + spontaneous. (*c*) Scatterplot of all stimulus conditions plotted against one another. Size and color of each point indicate density of points at that location. Examples of (*d*) low-flexibility and (*e*) high-flexibility modules. In *d* and *e*, modules have been recolored to match the nine-module partition detected using the spontaneous network alone. Nodes with gray colors have module assignments that cannot be easily mapped to one of the nine modules. (*f*) Similarity (z-score of the Rand index) of stimulus-evoked and spontaneous modular structure as a function of the inter-layer coupling parameter, *ω*. When *ω* is small, detected modules reflect specific stimuli; when *ω* is larger, detected modules are more similar to spontaneous modular structure. g) Node-level flexibility grouped according to modules. Note: high levels of heterogeneity across modules. (*h*) Node-level flexibility plotted in anatomical space. (*i*) Correlation of stimulus-evoked flexibility and participation coefficient estimated from spontaneous network data alone (*j*) Variation of flexibility patterns as the number of detected modules changes.

### Hub nodes reconfigure in response to stimuli

In the previous section we found that stimulus-evoked and spontaneous FC were highly correlated with one another across different stimulus conditions. In this section, we tease apart those differences. To address this question, we adapted a multi-layer modularity maximization procedure (see **Materials and Methods** for more details) [61], which has been used to study time-varying FC [62] and inter-individual differences at the macroscale [47]. This procedure entails treating each spontaneous and stimulus-evoked FC matrix as a “layer” in a multi-layer network ensemble. Layers are coupled to one another and the entire ensemble used as input for modularity maximization, thereby estimating modules in all layers simultaneously. As a result, this procedure allows us to effortlessly map modules across stimulus conditions and to identify flexible and inflexible nodes, i.e. those whose module assignments are consistently variable *versus* those that are stable.

The multi-layer modularity maximization framework includes an additional free parameter, *ω*, that can be tuned to different values so that the detected modules emphasize either the unique modular structure for each stimulus condition or the shared modular structure across conditions. When *ω* is large (emphasizing common modular structure), the detected multi-layer modules are highly similar to those detected in the previous section using the spontaneous networks, alone (Fig. 4d). This result is expected given the high correlation between stimulus-evoked and spontaneous FC. Here, we show as an example the module assignments that best match the nine-module partition shown in Figure 3g. When *ω* is set to smaller values, however, the algorithm detects modules that are uniquely suited to each layer and therefore more variable (Figure. 4e). We see this more clearly when we compare stimulus-evoked and spontaneous modular structure as a function of *ω* (Figure. 4f). The average similarity of stimulus-evoked and spontaneous modular structure, as measured by the z-score of the Rand index [63], increases monotonically as a function of *ω*.

Because the multi-layer modularity maximization framework preserves module labels across layers, it facilitates straightforward comparisons of modules from one stimulus condition with modules from another. This allows us to calculate each node’s “flexibility” [62, 64] a measure of its network variability across stimuli (see **Materials and Methods**). Intuitively, nodes with high flexibility are those that frequently switch module allegiance in response to a stimulus, whereas low-flexibility nodes are invariant and serve as stable anchors of network organization across different conditions. Here, we calculated the average flexibility for each node across all sets of multi-layer modules for which the mean number of modules was between 2 and 25. Interestingly, we found that flexible nodes were not uniformly distributed across modules, but were exhibited high levels of heterogeneity (Fig. 4f,g). Perhaps most surprising, when we compared participation coefficient (estimated from spontaneous FC alone) with flexibility, we found a strong positive correlation (*r* = 0.57; *p* < 0.05), indicating that spontaneous network structure may play a role in shaping network responses to stimuli.

Collectively, these findings suggest that spontaneous, intrinsic network architecture represents a powerful constraint on FC during stimulus-evoked conditions, both in terms of connectivity patterns as well as modular structure. More specifically, we find a common, stable modular core around which a flexible periphery of nodes reconfigure their connections as they adapt and respond to ongoing stimuli. More importantly, the nodes that appear most willing to reconfigure under task conditions are the same nodes that occupy positions of influence with respect to the spontaneous modular structure. These findings further validate observations made at the macroscale, wherein high participation areas overlap with poly-functional association cortex [24, 65, 66].

### Stability of network architecture across individuals

In the previous sections, we demonstrated the spontaneous FC is organized by geometry, exhibits hierarchical modules and hubs, and constrains stimulus-evoked FC. In this final section, we conclude by showing that the network structure of spontaneous FC appears to be conserved and similar across individual subjects. To compare network structure across subjects, we estimated spontaneous FC networks separately for subjects 8–18. Nodes and edges were defined exactly as before. In Fig. 5a-c we show example FC matrices for subjects 8, 11, and 17.

**FIG. 5.**
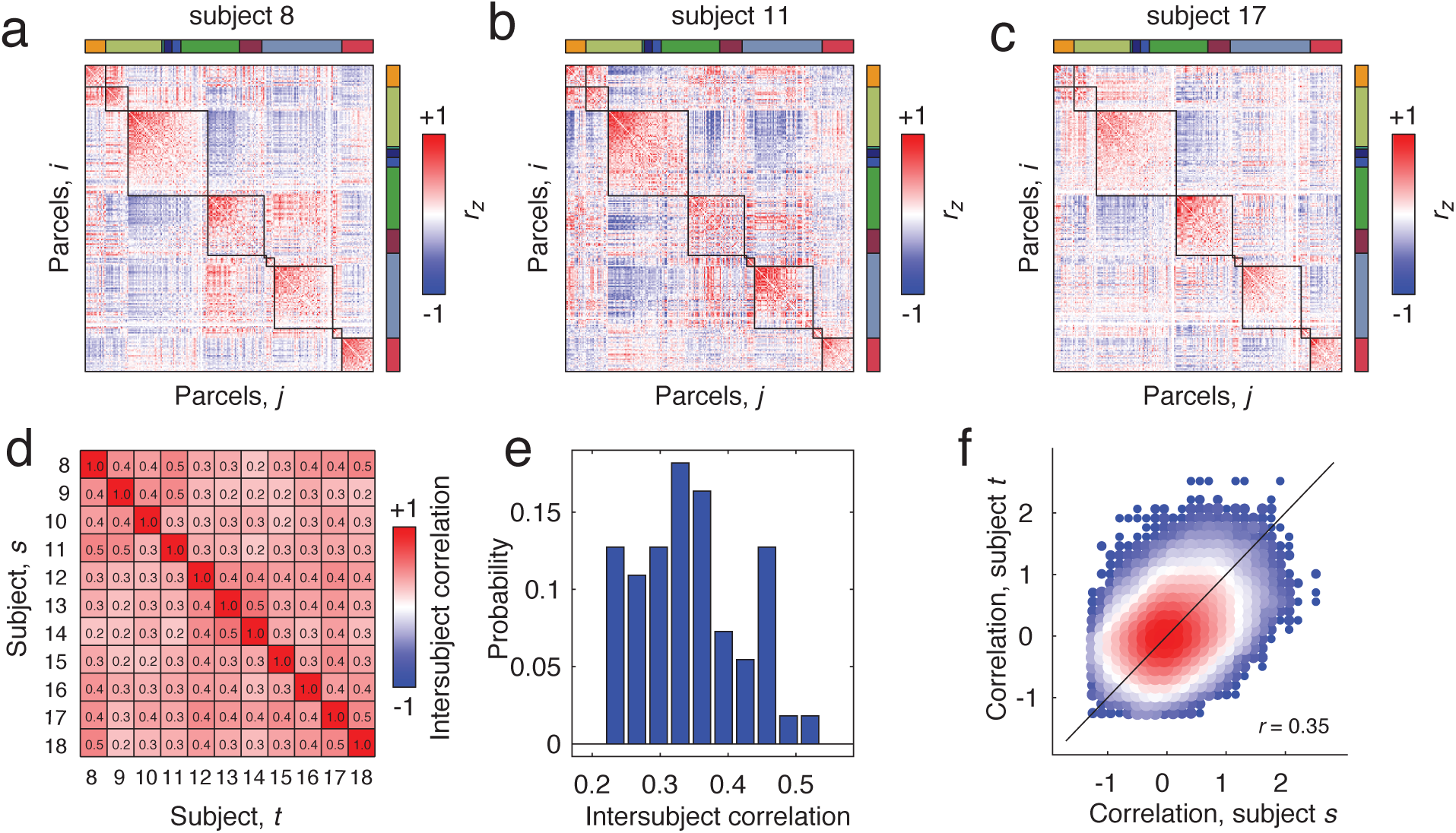
Intersubject similarity of spontaneous FC. Panels *a*, *b*, and *c* depict single-subject spontaneous FC matrices ordered according to the nine-module partition shown in Fig. 2g. (*d*) Correlation (similarity) matrix of upper triangle elements in subject-specific spontaneous FC matrices. (*e*) Distribution of elements in upper triangle of correlation matrix shown in panel *d*. (*f*) Scatterplot of functional connections for subjects *s* and *t*.

Upon visual inspection alone, the similarity of these matrices is apparent. Nonetheless, we quantified their similarity precisely by computing the correlation of their upper triangle elements with one another (Fig. 5d). We found that the mean correlation was *r* = 0.35 ± 0.08, indicating that while the correspondence of network architecture across subjects was imperfect, it was nonetheless robustly similar. We visualize these findings in two different way: In Fig. 5e we plot the distribution of inter-subject similarity and in Fig. 5f we show the two-dimensional histogram of connections plotted against one another.

In summary, these findings indicate that network structure of spontaneous FC is highly reproducible and conserved across individuals. Further, they suggest the presence of a neuro-functional blueprint shared across individuals. These findings contribute to the growing literature on uncovering sources of inter-subject variability in FC [67, 68] and extend this research domain from the macro-to the microscale.

## DISCUSSION

Nervous systems are fundamentally complex networks of interacting neurons, neuronal populations, and brain areas. Current research into the structure of these networks has begun to reveal basic organizing principles. Despite this, little is known about the architecture of biological neural networks at the microscale (cellular level). Here, we use network science methods to investigate the organization of microscale networks in larval zebrafish during spontaneous and stimulus conditions. Our analyses reveal the effect of geometry on network structure, a modular architecture and the hubs that span module boundaries, and the constraint of intrinsic network structure on stimulus-evoked activity. These features are analogous to those observed in whole-brain networks constructed at the macroscale; our findings and methods therefore serve as conceptual bridges, linking investigations of nervous system structure and functional across organizational scales. The work presented here serves as a methodological blueprint for future microscale network analyses and highlights several outstanding neuroscientific questions to be addressed with additional experiments and modeling.

### Bridging scales

Nervous systems exhibit meaningful organization and behavior across a wide range of spatial scales – from the level of cells and molecules up to brain areas [12]. However, different spatial scales are imaged using different technologies, resulting in data that is often modeled and analyzed using different statistical and mathematical methods. These distinctions in how spatial scales are researched can give rise to sometimes competing or contradictory accounts of nervous system organization and behavior; this is especially true when the neural phenomenon being studied is not clearly restricted to a single scale. Accordingly, there is a growing need for theoretical frameworks capable of explaining and modeling observations simultaneously at many scales.

In the present work, we use network science methodology to model and characterize whole-brain microscale networks. Though these methods have been widely used at the macroscale [8, 9, 69], they are only beginning to be applied to microscale datasets and have been restricted to networks reconstructions at the population level [34–36, 38, 70] or at the level of entire brains but without cellular resolution [51, 71, 72]. In extending the network approach to the whole-brain level and for networks reconstructed from single-cell observations, we provide a powerful demonstration of the utility of network science for gaining insight into nervous system architecture. Moreover, our work showcases network science as a framework with the ability to form bridges across spatial scales and, potentially, help reconcile disparate findings [7].

### Universal organizational principles

One of the goals of network neuroscience is to not simply *describe* a network, but to identify the principles by which nervous systems are organized [7]. In other words, what are the rules by which a network’s structure is shaped [44, 73–75] and are those rules universally true or do they apply only to particular organism and scale [30]? This goal is challenging to address, as most research that constitutes network neuroscience has focused on a single scale (macro) for a single organism (human), limiting the possibility of discovering scale- and species-invariant organizing principles.

Here, due to recent advances in cellular-level imaging [39, 41], we constructed whole-brain microscale networks and find, surprisingly, that these networks express analogous features at macroscale networks, suggesting that similar rules and principles are responsible. These features include connection weights with a strong geometric dependence, a balance between segregated and integrated brain function, and subtle stimulus-induced reconfigurations of network architecture. These findings indicate that networks reconstructed at the microscale may be organized according to analogous set of rules and principles, suggesting that these rules may be approximately scale-invariant. Moreover, these findings also suggests that, despite increases in dimensionality at the microscale, network organization can be succinctly described by a low-dimensional set of principles.

### Future directions

There are several ways that the work presented here could be extended in the future. First, although the network science methods we used provided novel insight into network structure at the microscale, those methods are relatively established in the network neuroscience literature [9, 56, 76]. Recently, increasingly novel methods have been developed that assay more nuanced aspects of network structure; in the long-term, it would be useful to test whether these new (but more complex) approaches based on topological data analysis [77], control theory [78], blockmodeling [79], and graph signal processing [80], can return give novel insights into microscale network structure and function. In the near-term, a potentially more fruitful approach could involve estimating time-varying FC to track fluctuations in network structure over short timescales and thereby gain insight into network dynamics [81].

Second, though networks analyzed here were reconstructed from single-cell recordings, we aggregated cells into parcels which we regarded as nodes. Though the parcels were defined to be both spatially and functionally homogeneous, each parcel-averaged fluorescence trace likely does not perfectly explain the variance of traces for every cell assigned to that parcel. In future work, it is critical to explore the effect of alternative parcellations on network statistics [82]. Additionally, it would be useful to explore parcellation-free approaches, wherein connections are estimated between individual cells (though this approach will likely scale poorly as number of recorded cells increases [83]).

### Limitations

Despite contributing to our understanding of the organizing principles underpinning microscale biological neural networks, this study has a number of important limitations. First, it is critical to note that our measure of connectivity is a linear correlation. Though this particular measure has been used extensively to model functional connectivity of slowly-evolving fMRI BOLD data [84] and performs well in recovering underlying network structure when applied to synthetic time series data [85], it is not a direct measure of structural connections (axonal projections), it is not equivalent to the coupling matrix in a dynamical systems model, does not indicate directedness of a connection, and is in no way a measure of causality or mechanism. Consequently, the results of all analyses are descriptive in nature. In future work, perturbational experiments and novel network reconstruction techniques will help clarify the causal role of network organization [86, 87].

Second, we infer connection weights based on zero-lag correlations of calcium fluorescence, which serves as an indirect measurement of a neuron’s activity. This indirectness coupled with lack of temporal precision (two volumes acquired per second), opens to the possibility that reconstructed networks are not fully recapitulating the “true” correlation structure of neurons’ activities. Advances in electrophysiological methods that enable recording from large number of neurons [88] and optical techniques for accelerating acquisition times [89] will prove useful in addressing these and related issues in future work.

## MATERIALS AND METHODS

### Data acquisition

Activity from a majority of neurons was recorded in larval zebrafish using light-sheet microscopy [41]. Activity was recorded under spontaneous conditions and also while subjects were presented with a series of visual stimuli. As reported in [90], calcium imaging data was recorded at ≈ 2 volumes/second for ≈ 50 minutes (or ≈ 6800 volumes). As in [41], the calcium indicator, GCaMP6f [91], was expressed pan-neuronally and fused to cell nuclei, allowing for the automatic segmented of cells [92]. The result was continuous fluorescence traces for ≈ 80, 000 cells per subject. As noted in [90], this number accounts for the majority of neurons in the brain, excluding extremely ventral areas. For complete details of data acquisition, see [90].

### Visual stimuli

In addition to spontaneous activity, subjects were imaging concurrent with the presentation of a series of visual stimuli (see [90] for complete details). Briefly, these stimuli included:

1. *Phototaxis*: subjects were presented with half-field black and white visual stimulus on either side. Presentations were followed by a whole-field white stimulus.
2. *Optomotor response*: Whole-field stripes moving in different directions.
3. *Looming response*: Expanding discs on either left or right side.
4. *Dark-flash response*: Sudden darkening of environment.

### Node definition

Rather than analyze networks where nodes correspond to individual cells, we focused on networks where the nodes represented clusters of cells, effectively reducing computational burden and facilitating straightforward interpretation. We developed a data-driven approach to assign cells to clusters that possessed three distinct properties: (1) spatial contiguity, (2) functional homogeneity, and (3) inter-subject consistency.

To generate clusters with this set of properties, we designed a multi-stage clustering algorithm. First, we aggregated normalized spatial coordinates of cells in the left hemisphere across all subjects and clustered them using k-means into *k* = 2500 contiguous spatial clusters (distance metric = Euclidean; number of replicates = 10). We then mirrored the centroids about the midline and assigned cells in the right hemisphere to the nearest spatial cluster centroid (5000 hemispherically symmetric clusters; mean spatial cluster size of 280 ± 73 cells). Because this procedure defines spatial clusters based on cells’ spatial locations, the resulting spatial clusters were also spatially contiguous. Moreover, because spatial clusters were defined using aggregated coordinates, spatial clusters contained cells from multiple subjects. Of the 5000 spatial clusters, 82.6% contained cells from all subjects and 92.5% contains cells from at least 75% of subjects.

Next, we concatenated spontaneous and stimulus-evoked activity and regressed the global signal from each cell’s fluorescence trace [93]. This procedure ensured that any observed fluctuations in fluorescence was not driven by changes in baseline fluorescence. Then for each spatial cluster and each subject, we extracted a cluster-averaged fluorescence trace, and computed pairwise correlations for all spatial clusters. We repeated this algorithm separately for both each hemisphere before averaging over subjects *and* hemispheres, resulting in a single 2500 × 2500 matrix of group- and hemisphere-averaged correlations (we were able to average over hemispheres because spatial clusters were defined to be hemispherically symmetric and so each spatial cluster had a homotopic partner). Next, we used this matrix to identify functional clusters with high average functional homogeneity by clustering its rows using k-means. We asked the algorithm to generate *k* = 100 functional clusters (Distance = correlation; number of replicates = 10). However, the resulting functional clusters were, in general, no longer spatially contiguous (i.e. functional clusters could include spatial clusters whose fluorescence traces were similar, but not necessarily proximal to one another). Accordingly, we extracted all spatial clusters assigned to each functional cluster and further divided them according to their spatial coordinates until the maximum diameter was < 200 pixels and each spatial cluster in a functional cluster was no fewer than 60 pixels from any other spatial cluster assigned to the same functional cluster. In the end, this procedure resulted in *N* = 256 *parcels*, each of which represented a node in our network.

### Network construction

To constuct networks, we mapped the *N* = 256 nodes defined in the previous section back to individual subjects and to single cells. For each subject and for each node, we extracted the average fluorescence trace its constituent cells and computed the pairwise correlation for all pairs of nodes. All correlation coefficients were subsequently Fisher transformed.

### Multi-resolution modularity maximization

Modularity maximization is a flexible, data-driven method for partitioning networks into sub-networks called “communities” or “modules” [76]. It operates according to a simple principle – compare observed connections against what would be expected under a null connectivity model. Modules, then, are groups of nodes whose observed internal density of connections is maximally greater than expected. This intuition is quantified using the modularity heuristic, *Q*, defined as:

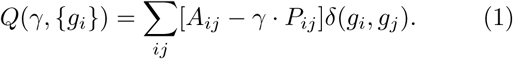

Here, *A_ij_* and *P_ij_* are the observed and expected connection weights between nodes *i* and *j*; *γ* is the structural resolution parameter; *g_i_* ∈ {1,…,*C*} is the module assignment of node *i*; and *δ*(*x*, *y*) is the Kronecker delta function, where *δ*(*x*, *y*) = 1 when *x* = *y* and is *δ*(*x*, *y*) = 0, otherwise. Effectively, this function calculates the total weight of within-module connections less that of the null model. Partitions that correspond to larger values of *Q* are considered to be, objectively, of higher quality.

#### Hierarchical modules

Here, we used two variants of modularity maximization to detect modules in zebrafish functional networks. The first seeks to relate coarse (large) and fine (small) modules to one another through a hierarchy [52] (http://github.com/LJeub/HierarchicalConsensus). This procedure takes advantage of the resolution parameter, *γ*, which can be tuned to detect modules of different sizes. Briefly, we sampled 10000 partitions at various values of *γ*, which results in modules of different sizes. We then summarized those modules using a co-assignment matrix whose elements indicated the fraction of all samples in which pairs of nodes were assigned to the same module. To generate a hierarchy, this co-assignment matrix was recursively clustered, with larger modules divided into smaller modules according to a statistical criterion. This procedure resulted is a series of hierarchically-related modules.

#### Multi-layer modules

We also used a second variant of modularity maximization, in which the *Q* heuristic is extended to “multi-layer” networks [61]. Here, layers represented connectivity matrices estimated under different stimulus conditions. For each layer, *s*, we calculated its corresponding modularity matrix, *B_s_* = *A_s_* −*γ*·*P_s_*. Node *i* in layer *s* was then linked to itself in all other layers by an inter-layer connection of weight *ω*. Next, modules were detected for all layers simultaneously by submitting the entire multi-layer ensemble – all layer-specific modularity matrices and inter-layer connections – to the modularity maximization algorithm (See Figure S2 for a schematic). The corresponding *Q* heuristic can be written:

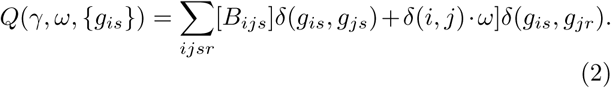

This multi-layer formulation has the distinct advantage over single-layer formulations, in that by detecting modules in all layers simultaneously, module labels are preserved across layers. That is, the observation of the same module label in layers *s* and *t* ≠ *s* indicates recurrence of a module. This fact facilitates straightforward module comparisons and enables the computation of a multi-layer flexibility measure, *f_i_*, whose value indicates the fraction of times that a node *i*’s module assignment differs across pairs of layers.

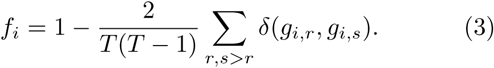

In the context of the current study, high-flexibility nodes

For both the hierarchical and multi-layer variants of modularity maximization, we used the generalized Louvain toolbox to optimize *Q* [94] (http://netwiki.amath.unc.edu). In both cases, we also used a uniform null connectivity model, i.e. *P_ij_* = 1, which helps mitigate issues related to the resolution limit [95] and is appropriate for use with correlation matrices [96].

#### Comparing modular structure

To compare two modular partitions, we used the z-score of the Rand index [63], a similarity measure that, intuitively, corrects the more common Rand index for the number and size of communities in the partitions. For two partitions, *X* and *Y*, we calculate their similarity as:

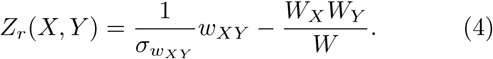

Here, *W* is the total number of node pairs in the network, *W_X_* and *W_Y_* are the number of pairs in the same modules in partitions *X* and *Y*, respectively, *w_XY_* is the number of pairs assigned to the same module in *both X* and *Y*, and 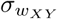 is the standard deviation of *w_XY_*. The value of *Z_r_*(*X*, *Y*) can be interpreted as how great, beyond chance, is the similarity of partitions *X* and *Y*.

### Defining network hubs

Hubs are considered nodes of disproportional importance to a network. We use three measures to identify hubs. The first measure is the flexibility metric. Nodes with high flexibility are those whose module affiliation changes across stimulus conditions and are therefore able to reconfigure in response to different stimuli or tasks.

The second metric used for hub classification is a node’s (weighted and signed) strength – i.e. the total weight of all of its connections:

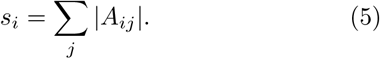

Intuitively, nodes with stronger connection weights, either positive or negative, may occupy positions of influence within the network.

Finally, we also defined hubs to be nodes with large participation coefficient [56]. A participation coefficient measures how uniformly a node’s connections (positive, in this case) are distributed across modules. A node that makes positive connections to many different modules will have a high participation coefficient, while a node whose connections are restricted to a small number of modules will have a low participation coefficient. More specifically, participation is calculated as:

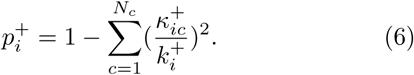

Here, *N_c_* is the number of modules, 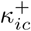 is the total weight of positive connections from node *i* to module *c*, and 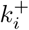 is the total weight of positive connections incident upon node *i*. Intuitively, nodes whose connections are distributed uniformly across modules have values of 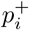 close to 1, while nodes whose connections are concentrated within a small number modules have values of 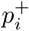 closer to 0. Participation coefficient was computed using functions from the Brain Connectivity Toolbox [9] (https://sites.google.com/site/bctnet/Home).

## DATA AVAILABILITY

All imaging data is available courtesy of M. Ahrens and X. Chen (https://github.com/xiuyechen/fishexplorer). All analysis scripts used in this manuscript are available from the authors upon request.

## ACKNOWLEGEMENTS

RFB is grateful to M. Ahrens and X. Chen for sharing zebrafish imaging data.

## AUTHOR CONTRIBUTIONS

RFB conceived of the project, performed all analyses, and wrote the paper.

---

**FIG. S1.**
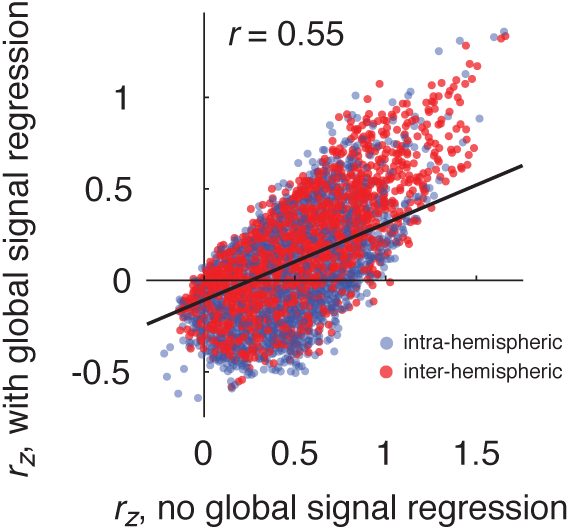
Comparison of spontaneous FC with and without global signal regression.

**FIG. S2.**
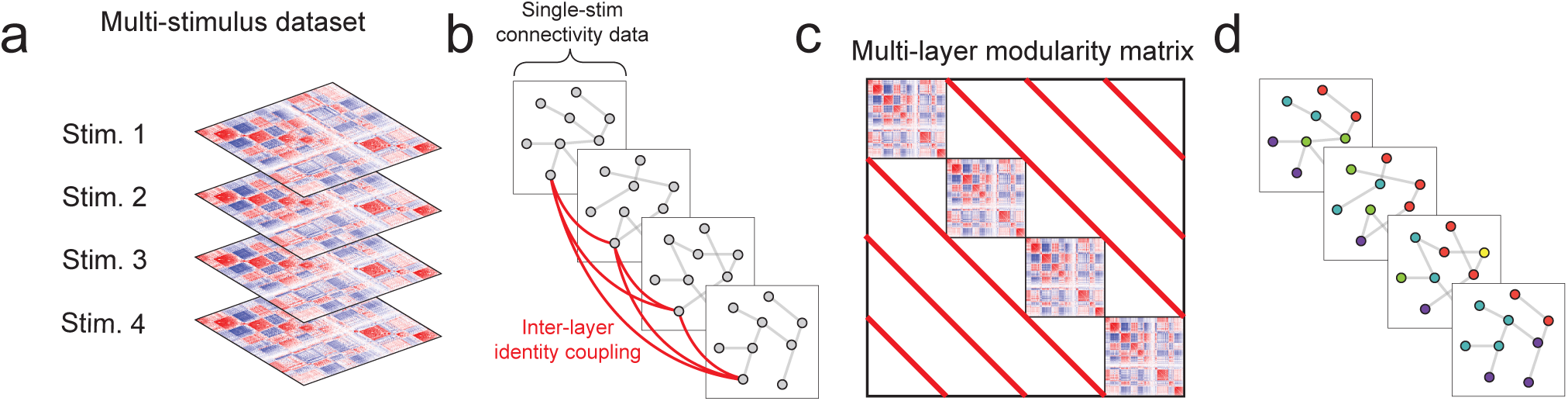
Multi-layer modularity maximization. We used multi-layer modularity maximization to simultaneously detect modules in spontaneous and stimulus-evoked FC. (*a*) The approach takes as input a series of single-layer connectivity matrices. In this case, the connectivity matrices represent FC estimated under different stimulus or spontaneous conditions. (*b*) Next, we add weak connections (of magnitude *ω*) from node *i* in layer *s* to itself in all other layers *t* ≠ *s*. (*c*) Next, we transform the connectivity matrices into single-layer modularity matrices by subtracted from each element the expected connection weight of connections (*γ*). These single-layer modularity matrices and the inter-layer connections are rearranged into a modularity tensor (shown here in planar form) which is submitted to the modularity maximization algorithm. (*d*) The multi-layer modularity tensor includes all layers (stimulus conditions) and estimates their modules simultaneously. The result is a set of modules labels that are conserved across layers. That is, a label *c* that appears in both layers *s* and *t* is interpreted as a recurrence of the same module. In summary, multi-layer modularity maximization effectively maps modules from one layer to another; this type of mapping, in general, is not possible using single-layer modularity maximization.

**FIG. S3.**
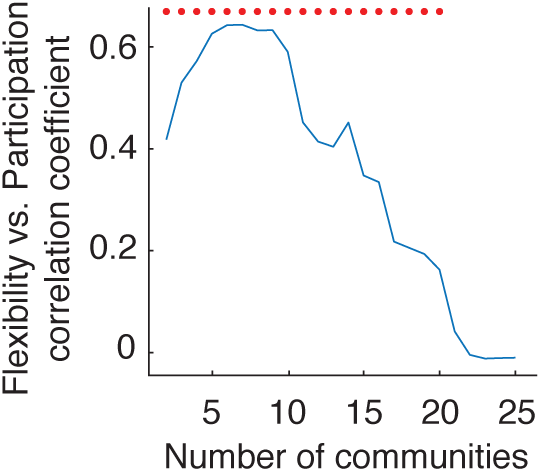
Multi-scale correspondence of participation and flexibility. In the main text, we show that a node’s average flexibility (over partitions of the network into different numbers of modules) is correlated with its average participation coefficient. Here, we show that this correspondence is not limited to a particular number of modules, but holds over a broad range. We plot the magnitude of the correlation between flexibility and participation coefficient as a function of the number of modules. Red stars indicate *p* < 0.05.

## References

[1] O. Sporns, G. Tononi, and R. Kötter, PLoS computational biology 1, e42 (2005).

[2] J. Downar, A. P. Crawley, D. J. Mikulis, and K. D. Davis, Nature neuroscience 3, 277 (2000).

[3] H. Ko, S. B. Hofer, B. Pichler, K. A. Buchanan, P. J. Sjöström, and T. D. Mrsic-Flogel, Nature 473, 87 (2011).

[4] T. Hartley, C. Lever, N. Burgess, and J. O’Keefe, Phil. Trans. R. Soc. B 369, 20120510 (2014).

[5] J. Jacobs, C. T. Weidemann, J. F. Miller, A. Solway, J. F. Burke, X.-X. Wei, N. Suthana, M. R. Sperling, A. D. Sharan, I. Fried, et al., Nature neuroscience 16, 1188 (2013).

[6] R. N. Spreng, W. D. Stevens, J. P. Chamberlain, A. W. Gilmore, and D. L. Schacter, Neuroimage 53, 303 (2010).

[7] D. S. Bassett and O. Sporns, Nature neuroscience 20, 353 (2017).

[8] E. Bullmore and O. Sporns, Nature Reviews Neuroscience 10, 186 (2009).

[9] M. Rubinov and O. Sporns, Neuroimage 52, 1059 (2010).

[10] H.-J. Park and K. Friston, Science 342, 1238411 (2013).

[11] G. Deco, V. K. Jirsa, P. A. Robinson, M. Breakspear, and K. Friston, PLoS computational biology 4, e1000092 (2008).

[12] R. F. Betzel and D. S. Bassett, Neuroimage 160, 73 (2017).

[13] M. Schröter, O. Paulsen, and E. T. Bullmore, Nature Reviews Neuroscience 18, 131 (2017).

[14] R. C. Craddock, S. Jbabdi, C.-G. Yan, J. T. Vogelstein, F. X. Castellanos, A. Di Martino, C. Kelly, K. Heberlein, S. Colcombe, and M. P. Milham, Nature methods 10, 524 (2013).

[15] A. Fornito, A. Zalesky, and M. Breakspear, Nature Reviews Neuroscience 16, 159 (2015).

[16] A. Di Martino, D. A. Fair, C. Kelly, T. D. Satterthwaite, F. X. Castellanos, M. E. Thomason, R. C. Craddock, B. Luna, B. L. Leventhal, X.-N. Zuo, et al., Neuron 83, 1335 (2014).

[17] M. Rubinov, Nature communications 7, 13812 (2016).

[18] R. F. Betzel, A. Avena-Koenigsberger, J. Goñi, Y. He, M. A. De Reus, A. Griffa, P. E. Vértes, B. Mišic, J.-P. Thiran, P. Hagmann, et al., Neuroimage 124, 1054 (2016).

[19] O. Sporns, G. Tononi, and G. Edelman, Behavioural brain research 135, 69 (2002).

[20] J. R. Cohen and M. D’Esposito, Journal of Neuroscience 36, 12083 (2016).

[21] O. Sporns and R. F. Betzel, Annual review of psychology 67, 613 (2016).

[22] D. Meunier, R. Lambiotte, A. Fornito, K. Ersche, and E. T. Bullmore, Frontiers in neuroinformatics 3, 37 (2009).

[23] M. P. van den Heuvel and O. Sporns, Trends in cognitive sciences 17, 683 (2013).

[24] J. D. Power, B. L. Schlaggar, C. N. Lessov-Schlaggar, and S. E. Petersen, Neuron 79, 798 (2013).

[25] E. Bullmore and O. Sporns, Nature Reviews Neuroscience 13, 336 (2012).

[26] B. L. Chen, D. H. Hall, and D. B. Chklovskii, Proceedings of the National Academy of Sciences 103, 4723 (2006).

[27] R. F. Betzel and D. S. Bassett, Proceedings of the National Academy of Sciences, 201720186 (2018).

[28] M. W. Cole, D. S. Bassett, J. D. Power, T. S. Braver, and S. E. Petersen, Neuron 83, 238 (2014).

[29] D. H. Schultz and M. W. Cole, Journal of Neuroscience 36, 8551 (2016).

[30] M. P. Van den Heuvel, E. T. Bullmore, and O. Sporns, Trends in cognitive sciences 20, 345 (2016).

[31] M. Rubinov, R. J. Ypma, C. Watson, and E. T. Bullmore, Proceedings of the National Academy of Sciences 112, 10032 (2015).

[32] M. D. Humphries, Network Neuroscience 1, 324 (2017).

[33] B. Dann, J. A. Michaels, S. Schaffelhofer, and H. Scherberger, Elife 5, e15719 (2016).

[34] J. G. Orlandi, J. Soriano, E. Alvarez-Lacalle, S. Teller, and J. Casademunt, Nature Physics 9, 582 (2013).

[35] S. P. Faber, N. M. Timme, J. M. Beggs, and E. L. Newman, Network Neuroscience, 1 (2018).

[36] H. Yamamoto, S. Moriya, K. Ide, T. Hayakawa, H. Akima, S. Sato, S. Kubota, T. Tanii, M. Niwano, S. Teller, et al., Science advances 4, eaau4914 (2018).

[37] W.-C. A. Lee, V. Bonin, M. Reed, B. J. Graham, G. Hood, K. Glattfelder, and R. C. Reid, Nature 532, 370 (2016).

[38] A. M. Bruno, W. N. Frost, and M. D. Humphries, Neuron 86, 304 (2015).

[39] M. B. Ahrens, J. M. Li, M. B. Orger, D. N. Robson, A. F. Schier, F. Engert, and R. Portugues, Nature 485, 471 (2012).

[40] M. B. Ahrens, M. B. Orger, D. N. Robson, J. M. Li, and P. J. Keller, Nature methods 10, 413 (2013).

[41] N. Vladimirov, Y. Mu, T. Kawashima, D. V. Bennett, C.-T. Yang, L. L. Looger, P. J. Keller, J. Freeman, and M. B. Ahrens, Nature methods 11, 883 (2014).

[42] P. J. Keller and M. B. Ahrens, Neuron 85, 462 (2015).

[43] P. Bellec, V. Perlbarg, S. Jbabdi, M. Pélégrini-Issac, J. L. Anton, J. Doyon, and H. Benali, Neuroimage 29, 1231 (2006).

[44] P. E. Vértes, A. F. Alexander-Bloch, N. Gogtay, J. N. Giedd, J. L. Rapoport, and E. T. Bullmore, Proceedings of the National Academy of Sciences, 201111738 (2012).

[45] J. Stiso and D. Bassett, arXiv preprint arXiv:1807.04691 (2018).

[46] S. B. Laughlin and T. J. Sejnowski, Science 301, 1870 (2003).

[47] R. F. Betzel, M. A. Bertolero, E. M. Gordon, C. Gratton, N. U. Dosenbach, and D. S. Bassett, bioRxiv, 413278 (2018).

[48] R. F. Betzel, A. Griffa, A. Avena-Koenigsberger, J. Goñi, J.-P. Thiran, P. Hagmann, and O. Sporns, Network Science 1, 353 (2013).

[49] J. D. Power, A. L. Cohen, S. M. Nelson, G. S. Wig, K. A. Barnes, J. A. Church, A. C. Vogel, T. O. Laumann, F. M. Miezin, B. L. Schlaggar, et al., Neuron 72, 665 (2011).

[50] M. A. Bertolero, B. T. Yeo, and M. D’Esposito, Proceedings of the National Academy of Sciences 112, E6798 (2015).

[51] M. P. Vanni, A. W. Chan, M. Balbi, G. Silasi, and T. H. Murphy, Journal of Neuroscience, 3560 (2017).

[52] L. G. Jeub, O. Sporns, and S. Fortunato, Scientific reports 8, 3259 (2018).

[53] P. N. Taylor, Y. Wang, and M. Kaiser, Scientific reports 7, 39859 (2017).

[54] O. Sporns, Current opinion in neurobiology 23, 162 (2013).

[55] G. Deco, G. Tononi, M. Boly, and M. L. Kringelbach, Nature Reviews Neuroscience 16, 430 (2015).

[56] R. Guimera and L. A. N. Amaral, nature 433, 895 (2005).

[57] M. A. Bertolero, B. Yeo, D. S. Bassett, and M. D’Esposito, arXiv preprint arXiv:1803.08109 (2018).

[58] C. Honey, O. Sporns, L. Cammoun, X. Gigandet, J.-P. Thiran, R. Meuli, and P. Hagmann, Proceedings of the National Academy of Sciences 106, 2035 (2009).

[59] J. Goñi, M. P. van den Heuvel, A. Avena-Koenigsberger, N. V. de Mendizabal, R. F. Betzel, A. Griffa, P. Hagmann, B. Corominas-Murtra, J.-P. Thiran, and O. Sporns, Proceedings of the National Academy of Sciences 111, 833 (2014).

[60] R. F. Betzel, M. A. Bertolero, and D. S. Bassett, bioRxiv, 355016 (2018).

[61] P. J. Mucha, T. Richardson, K. Macon, M. A. Porter, and J.-P. Onnela, science 328, 876 (2010).

[62] D. S. Bassett, N. F. Wymbs, M. A. Porter, P. J. Mucha, J. M. Carlson, and S. T. Grafton, Proceedings of the National Academy of Sciences (2011).

[63] A. L. Traud, E. D. Kelsic, P. J. Mucha, and M. A. Porter, SIAM review 53, 526 (2011).

[64] D. S. Bassett, M. A. Porter, N. F. Wymbs, S. T. Grafton, J. M. Carlson, and P. J. Mucha, Chaos: An Interdisciplinary Journal of Nonlinear Science 23, 013142 (2013).

[65] M. W. Cole, J. R. Reynolds, J. D. Power, G. Repovs, A. Anticevic, and T. S. Braver, Nature neuroscience 16, 1348 (2013).

[66] T. P. Zanto and A. Gazzaley, Trends in cognitive sciences 17, 602 (2013).

[67] M. L. Seghier and C. J. Price, Trends in cognitive sciences (2018).

[68] E. M. Gordon, T. O. Laumann, A. W. Gilmore, D. J. Newbold, D. J. Greene, J. J. Berg, M. Ortega, C. Hoyt-Drazen, C. Gratton, H. Sun, et al., Neuron 95, 791 (2017).

[69] Y. He and A. Evans, Current opinion in neurology 23, 341 (2010).

[70] S. F. Muldoon, I. Soltesz, and R. Cossart, Proceedings of the National Academy of Sciences 110, 3567 (2013).

[71] E. M. Lake, X. Ge, X. Shen, P. Herman, F. Hyder, J. A. Cardin, M. J. Higley, D. Scheinost, X. Papademetris, M. C. Crair, et al., bioRxiv, 464305 (2018).

[72] D. Barson, A. S. Hamodi, X. Shen, G. Lur, R. T. Constable, J. Cardin, M. Crair, and M. Higley, bioRxiv, 468348 (2018).

[73] M. Ercsey-Ravasz, N. T. Markov, C. Lamy, D. C. Van Essen, K. Knoblauch, Z. Toroczkai, and H. Kennedy, Neuron 80, 184 (2013).

[74] R. F. Betzel and D. S. Bassett, Journal of The Royal Society Interface 14, 20170623 (2017).

[75] A. Goulas, R. F. Betzel, and C. C. Hilgetag, bioRxiv, 385369 (2018).

[76] M. E. Newman and M. Girvan, Physical review E 69, 026113 (2004).

[77] M. Saggar, O. Sporns, J. Gonzalez-Castillo, P. A. Bandettini, G. Carlsson, G. Glover, and A. L. Reiss, Nature communications 9, 1399 (2018).

[78] S. Gu, F. Pasqualetti, M. Cieslak, Q. K. Telesford, B. Y. Alfred, A. E. Kahn, J. D. Medaglia, J. M. Vettel, M. B. Miller, S. T. Grafton, et al., Nature communications 6, 8414 (2015).

[79] R. F. Betzel, J. D. Medaglia, and D. S. Bassett, Nature Communications 9, 346 (2018).

[80] W. Huang, T. A. Bolton, J. D. Medaglia, D. S. Bassett, A. Ribeiro, and D. Van De Ville, Proceedings of the IEEE (2018).

[81] R. M. Hutchison, T. Womelsdorf, E. A. Allen, P. A. Bandettini, V. D. Calhoun, M. Corbetta, S. Della Penna, J. H. Duyn, G. H. Glover, J. Gonzalez-Castillo, et al., Neuroimage 80, 360 (2013).

[82] J. Wang, L. Wang, Y. Zang, H. Yang, H. Tang, Q. Gong, Z. Chen, C. Zhu, and Y. He, Human brain mapping 30, 1511 (2009).

[83] T. H. Kim, Y. Zhang, J. Lecoq, J. C. Jung, J. Li, H. Zeng, C. M. Niell, and M. J. Schnitzer, Cell reports 17, 3385 (2016).

[84] O. David, D. Cosmelli, and K. J. Friston, Neuroimage 21, 659 (2004).

[85] S. M. Smith, K. L. Miller, G. Salimi-Khorshidi, M. Webster, C. F. Beckmann, T. E. Nichols, J. D. Ramsey, and M. W. Woolrich, Neuroimage 54, 875 (2011).

[86] L. Huang, J. Kebschull, D. Furth, S. Musall, M. T. Kaufman, A. K. Churchland, and A. Zador, bioRxiv, 422477 (2018).

[87] B. Lansdell and K. Kording, bioRxiv, 253351 (2018).

[88] J. J. Jun, N. A. Steinmetz, J. H. Siegle, D. J. Denman, M. Bauza, B. Barbarits, A. K. Lee, C. A. Anastassiou, A. Andrei, C¸. Aydın, et al., Nature 551, 232 (2017).

[89] A. J. Sadovsky, P. B. Kruskal, J. M. Kimmel, J. Ostmeyer, F. B. Neubauer, and J. N. MacLean, Journal of neurophysiology 106, 1591 (2011).

[90] X. Chen, Y. Mu, Y. Hu, A. T. Kuan, M. Nikitchenko, O. Randlett, H. Sompolinsky, F. Engert, and M. B. Ahrens, bioRxiv, 289413 (2018).

[91] T.-W. Chen, T. J. Wardill, Y. Sun, S. R. Pulver, S. L. Renninger, A. Baohan, E. R. Schreiter, R. A. Kerr, M. B. Orger, V. Jayaraman, et al., Nature 499, 295 (2013).

[92] T. Kawashima, M. F. Zwart, C.-T. Yang, B. D. Mensh, and M. B. Ahrens, Cell 167, 933 (2016).

[93] J. D. Power, A. Mitra, T. O. Laumann, A. Z. Snyder, B. L. Schlaggar, and S. E. Petersen, Neuroimage 84, 320 (2014).

[94] I. S. Jutla, L. G. Jeub, and P. J. Mucha, URL http://netwiki.amath.unc.edu/GenLouvain (2011).

[95] V. A. Traag, P. Van Dooren, and Y. Nesterov, Physical Review E 84, 016114 (2011).

[96] M. Bazzi, M. A. Porter, S. Williams, M. McDonald, D. J. Fenn, and S. D. Howison, Multiscale Modeling & Simulation 14, 1 (2016).

